# Evaluation of purine-nucleoside degrading ability and *in vivo* uric acid lowering of *Streptococcus thermophilus* (ID-PDP3), a novel antiuricemia strain

**DOI:** 10.1101/2023.10.12.562018

**Authors:** Dayoung Kim, Jin Seok Moon, Ji Eun Kim, Ye-Ji Jang, Han Sol Choi, Ikhoon Oh

## Abstract

This study evaluated 15 lactic acid bacteria in terms of their ability to degrade inosine and hypoxanthine—which are the intermediates in purine metabolism—for the management of hyperuricemia and gout. After a preliminary screening based on HPLC, CR1 (*Lactiplantibacillus plantarum*) and GZ1 (*Lactiplantibacillus pentosus*) showed the highest nucleoside degrading rates and were therefore selected for further characterization. IDCC 2201 (*S. thermophilus*), which possessed the *hpt* gene encoding hypoxanthine-guanine phosphoribosyltransferase (HGPRT) and exhibited purine degradation, was also selected. These three selected strains were examined in terms of the effect of probiotics on lowering serum uric acid in a rat model of potassium oxonate (PO)-induced hyperuricemia. Among them, the level of serum uric acid was most reduced by IDCC 2201 (p < 0.05). Further, analysis of the microbiome showed that administration of IDCC 2201 recovered the ratio of Bacteroidetes/Firmicutes in the intestinal microbial composition unbalanced by hyperuricemia and showed a difference in the intestinal microbial composition compared to the group administered with allopurinol. Moreover, intestinal short-chain fatty acids (SCFAs) were significantly increased. Ultimately, the findings show that IDCC 2201 lowers uric acid levels by degrading purine-nucleosides and restores intestinal flora and SCFAs, ultimately suggesting that IDCC 2201 is a promising candidate for use as an adjuvant treatment in patients with hyperuricemia.

## Introduction

Uric acid is an end product of the metabolism of purines in the body, and elevated serum uric acid levels have been shown to cause both hyperuricemia and gout [1,2]. The typical range of uric acid in human plasma is from 4 to 7 mg/dL, and concentrations outside this range are considered to be abnormal and are referred to as hyperuricemia or hypouricemia, respectively [3]. Long-term excess uric acid at levels above 7 mg/dL tends to lead to the formation of crystalline uric acid deposits on joint and cartilage surfaces. The accumulation of uric acid crystals in joints and soft tissues can cause inflammation and excruciating pain, and this is clinically diagnosed as gout. Although humans do not produce enzymes that degrade uric acid, some bacteria in the intestine can degrade one-third of dietary and endogenous uric acid, so recent hyperuricemia-related studies have focused on analyzing intestinal flora [4–6]. The beneficial effects of probiotics include maintaining gut microbiome composition, regulating immune response, managing metabolic disorders, etc. A number of studies have shown that these beneficial effects are related to the strain itself and the regulation of the gut microbiome [7]. Studies have shown that *Lactobacillus gasseri* PA-3 can lower serum uric acid levels by reducing the intestinal absorption of both inosine and adenosine-related compounds [8]. In one clinical trial, 25 patients with hyperuricemia and gout were given yogurt containing PA-3, and the results showed that PA-3 improved serum uric acid levels in the patients [9]. *Lactiplantibacillus plantarum* DM9218-A strain isolated from food significantly reduced serum uric acid levels in rats with induced hyperuricemia, and it also showed a preventive effect against hyperuricemia, suggesting its potential use as an adjuvant treatment for hyperuricemia [10]. Another study demonstrated the ability of *Lactobacillus paracasei* S12 strain isolated from pickles to degrade uric acid *in vitro* and *in vivo*, thus revealing its potential to reduce kidney damage [11]. It has also been reported that *Lactobacillus fermentum* 9-4 strain, which has been confirmed to have the ability to degrade purine-nucleosides *in vitro*, lowers uric acid by degrading inosine and guanosine [12]. *Lactiplantibacillus pentosus* P2020 has been reported to reduce renal inflammation by inhibiting inflammation-related signals and to reduce uric acid levels by affecting the expression of proteins involved in uric acid transport [13]. Altogether, these studies demonstrate that probiotics hold promise as an alternative adjuvant treatment for hyperuricemia to improve renal and intestinal function [13,14].

In this study, we screened and isolated a candidate strain with high purine-degrading activity, and we assessed the anti-hyperuricemia activity in potassium oxonate (PO)-induced hyperuricemia mice. We also investigated a possible mechanism for lowering the serum uric acid level through fecal analysis.

## Materials and methods

### Isolation, DNA extraction, and identification of strains

In this study, five type strains and 10 strains isolated from food were used to screen lactic acid bacteria with high purine decomposition activity (Table 1). To this end, *Lacticaseibacillus*, *Lactiplantibacillus*, *Lactobacillus*, *Limosilactobacillus*, and *Streptococcus* were cultured in Lactobacilli MRS broth (BD, MD, USA). All strains were incubated at 37 °C for 24 hours.

**Table 1.**
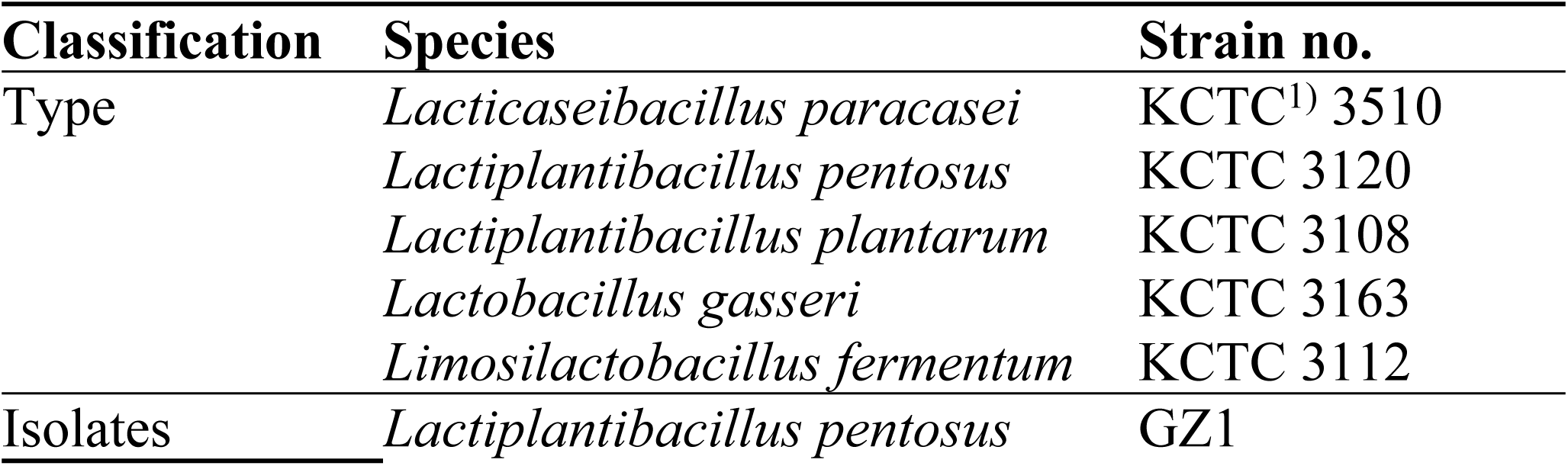
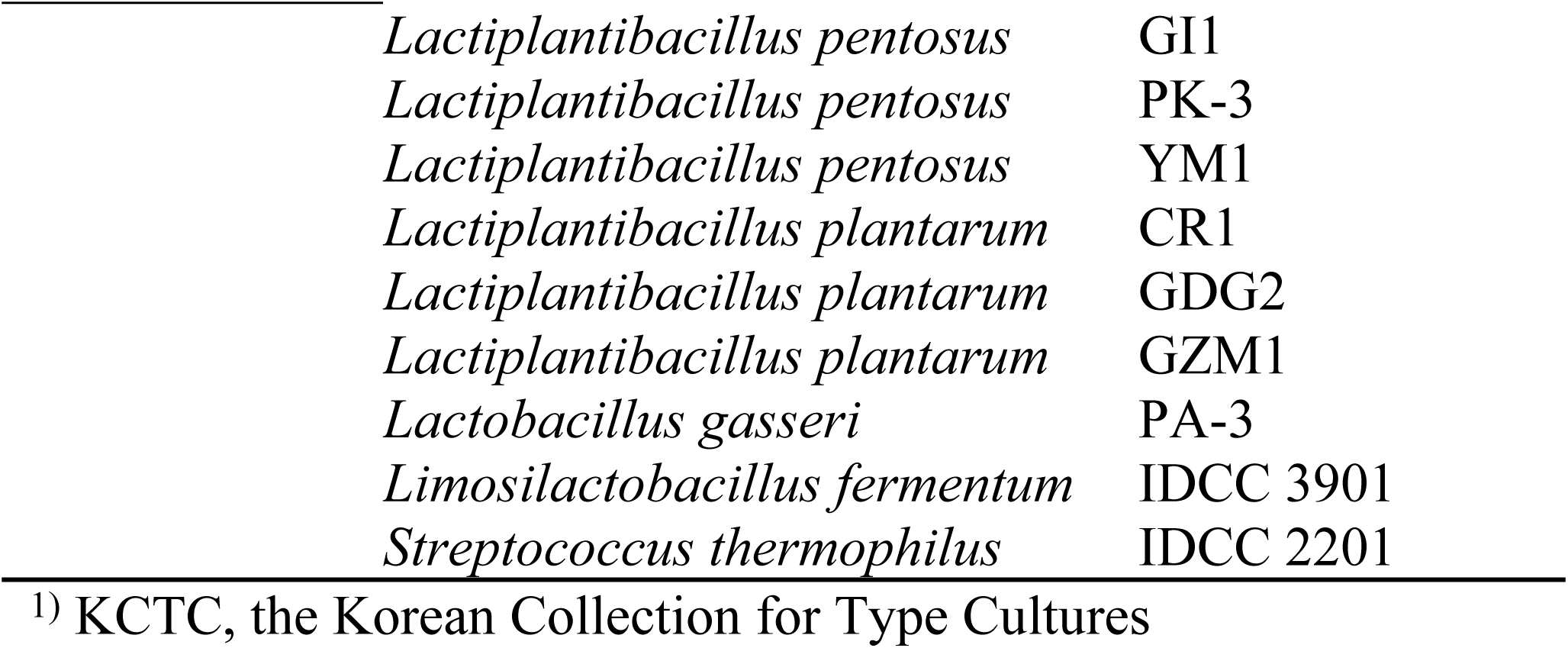
List of strains used in this study.

For the extraction of genomic DNA, cell cultures were centrifuged at 13,000 g for 10 minutes, and cell pellets were extracted using the DNeasy Blood and Tissue kit (Qiagen, Hilden, Germany) according to the manufacturer’s recommendations. Genomic DNA concentration was calculated using a DS-11+ spectrophotometer (DeNovix, DE, USA) and stored at −20 °C until being used as a template.

The extracted genomic DNAs were amplified with a 785F/907R primer pair and a 27F/1492R primer pair, and the strains were identified via 16S rRNA gene sequencing.

### Screening of LABs capable of degrading inosine and hypoxanthine

To evaluate the inosine and hypoxanthine assimilating ability, LAB strain was inoculated in MRS and cultured for 24 h at 37 °C under anaerobic conditions. To begin, 5 mL of the culture broth was centrifuged at 4,000 g, 4 °C for 10 minutes. The cells were then washed twice with 3 mL saline, suspended with 1.25 mM of inosine-hypoxanthine solution in potassium phosphate buffer (0.1 M, pH 7.0), and incubated at 37 °C for 60 minutes with shaking at 120 rpm. Then, the solution was centrifuged at 4,000 g, 4 °C for 10 minutes. After filtering using a 0.45-μm Millipore filter (Whatman, Maidstone, England), the supernatant was injected into the HPLC device.

Inosine-hypoxanthine solution was measured using an HPLC-UV detector according to the protocol described by Law et al. [15]. The prepared supernatant was analyzed using a Waters T3 column (5 um, 250 × 4.6 mm) connected to an HPLC pump (Alliance e2695; Waters Corp., Massachusetts, USA) and a Waters 2998 UV detector. The column temperature was maintained at 25 °C; mobile phase A was 0.1% phosphoric acid in water and mobile phase B was acetonitrile. The gradient program was as follows: 0–10 min, 98% A isocratic; 10–15 min, 98% A to 85% A; 15–25 min, 85% A to 20% A; 25–30 min, and 98% A isocratic at a flow rate of 0.8 mL/min. Inosine and hypoxanthine were detected at 254 nm with the retention times of 21.8 and 11.8 minutes and quantified through comparison with the standard.

The quantification of each compound was performed based on the peak area value of the HPLC chart. The resolution of inosine-hypoxanthine was measured and calculated while following a previously described research method [16]. The formula for calculating the degradation rate is as follows: degradation rate (%) = 100 - ((peak area at 1 hour / peak area at 0 hour) × 100).

### Purine metabolism related gene

The enzyme hypoxanthine-guanine phosphoribosyltransferase (HGPRT), which lowers uric acid levels by converting hypoxanthine and guanine to guanosine monophosphate (GMP) and inosine monophosphate (IMP), was selected as an important factor [17] (Fig 1). The *hpt* gene encoding this enzyme was searched through the United Protein Database (UniProt: https://www.uniprot.org) database.

**Fig 1.**
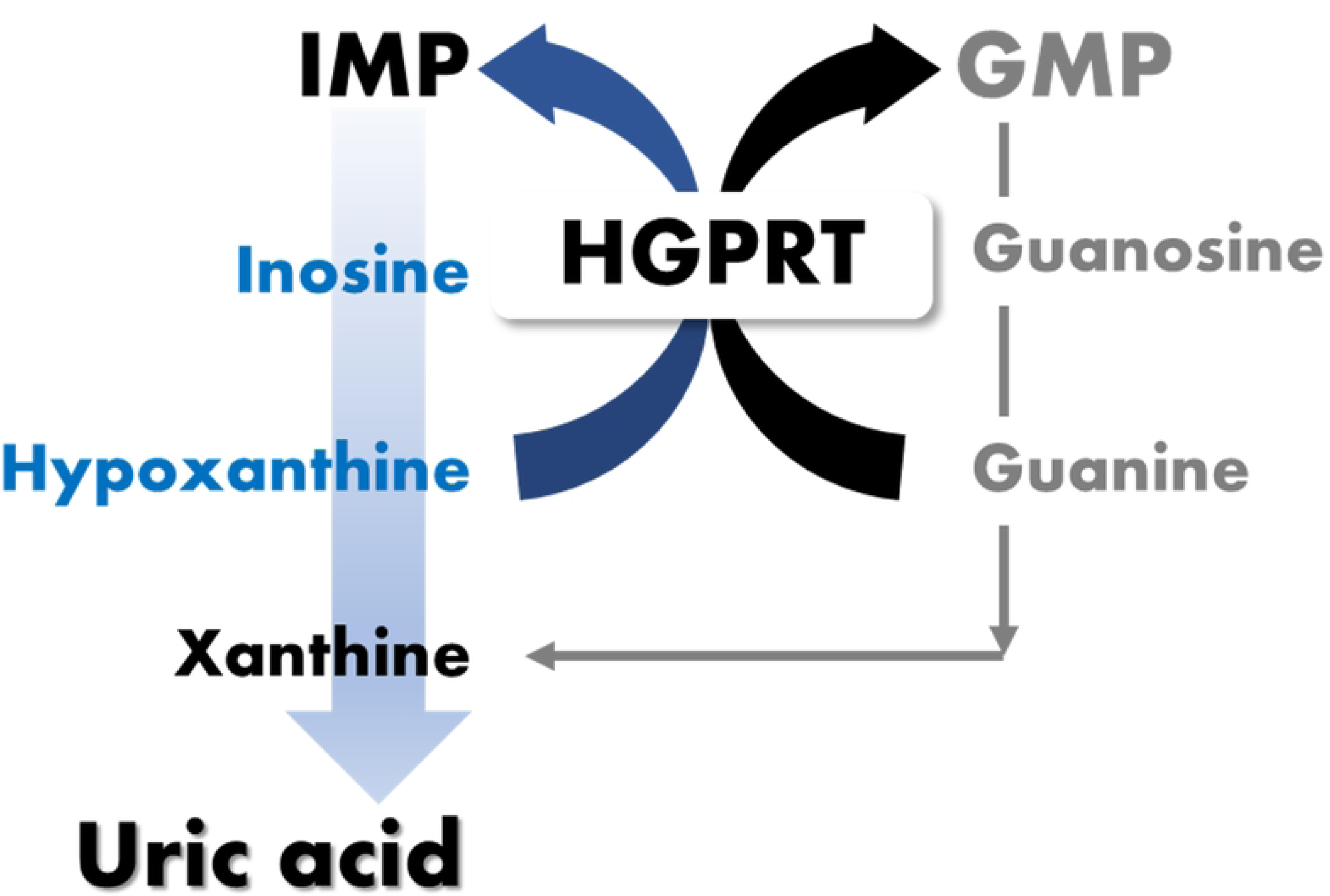
Overview of purine metabolism by HGPRT. The purine metabolic system represents a pathway by which hypoxanthine and guanine can be salvaged by HGPRT. HGPRT serves to catalyze the salvage synthesis of IMP and GMP from the purine bases hypoxanthine and guanine. HGPRT prevents the accumulation of substrates that are converted to uric acid.

### Experimental design of candidate strains of probiotics

The animal experiment in this study was performed at the Dt&CRO Efficacy Evaluation Center (Yongin, Republic of Korea) with approval from the Institutional Animal Care and Use Committee (IACUC approval number: 210317). To this end, 54 8-week-old male Sprague-Dawley (SD) rats obtained from Orientbio Inc. (Seongnam, Republic of Korea) were adapted to the environment at 22 ± 3 °C and 50 ± 20% humidity for 7 days with a 12-hour light/dark cycle. After 7 days of acclimatization, rats were randomly assigned to nine groups with six rats in each group. The nine groups were as follows: (a) normal group (G1); (b) hyperuricemia group (G2); (c) CR1 (1×109 CFU/ day, G3); (d) GZ1 (1×109 CFU/ day, G4); (e) IDCC 2201 (1×109 CFU/ day, G5); (f) CR1+IDCC 2201 (1×109 CFU/ day, G6); (g) GZ1+IDCC 2201 (1×109 CFU/ day, G7); (h) PA-3 (1×109 CFU/ day, G8); and (i) Allopurinol (50 mg/kg rat/day, G9). The normal group was intraperitoneally administered a dose of 8 mL/kg of 0.5% carboxy methyl cellulose (Sigma-Aldrich, MO, USA) as an excipient. The hyperuricemia induction and test substance administration groups were intraperitoneally administered a dose of PO (TCI, Tokyo, Japan) 8 mL/kg (250 mg/kg) for 7 days. Test materials were administered 1 hour after PO administration. On day 7, before animal sacrifice, blood was collected from the tail vein, and biochemical indicators such as blood uric acid were measured using a blood biochemical analyzer (Hitachi Ltd., Tokyo, Japan). For intestinal microflora analysis, fecal samples were collected from the animals on the day of test termination and stored at −80 °C.

### Experimental design of IDCC 2201 by concentration

The IDCC 2201 concentration-dependent animal experiment was performed in the same manner as described in ‘Experimental design of candidate strains of probiotics’. The animal experiment was approved by IACUC (approval number: 220203) and conducted at the Dt&CRO Efficacy Evaluation Center. For this experiment, 45 8-week-old male SD rats were obtained and randomly assigned to five groups with eight rats in each group. The test substance administration group was given concentrations of IDCC 2201 1×109 CFU/day, 1×108 CFU/day, and 1×107 CFU/day, respectively. The groups were as follows: (a) normal group (G1); (b) hyperuricemia group (G2); (c) IDCC 2201-SH (1×109 CFU/day, G3); (d) IDCC 2201-SM (1×108 CFU/day, G4); (e) IDCC 2201-SL (1×107 CFU/day, G5).

### Fecal samples analysis

#### Fecal DNA purification and metagenome sequencing for gut microbiome analysis

For metagenome analysis, fecal DNA was extracted with the ExgeneTM Stool DNA Mini Kit (GeneAll Biotechnology, Seoul, Republic of Korea) according to the manufacturer’s protocol. The V3-V4 region of the bacterial 16S rRNA gene was amplified using barcoded universal primers, 341F and 805R. Microbiome profiling was conducted on the 16S-based Microbial Taxonomic Profiling platform provided by EzBioCloud Apps (ChunLab Inc., Seoul, Republic of Korea).

#### HPLC analysis of short-chain fatty acids (SCFAs) in fecal samples

SCFAs in fecal samples were quantified using HPLC analysis [18]. To extract SCFAs in the fecal samples, 1.0 g of each fecal sample was thawed, suspended in 8 mL of water, and homogenized using a vortex mixer for 5 minutes. Next, the solution was centrifuged at 9500 rpm for 10 minutes at 4 °C. The resulting supernatant was then filtered through a 0.2-μm cellulose acetate/surfactant-free membrane filter, and 10 μL was injected into a Waters e2695 HPLC system equipped with a Waters 2998 UV detector. Afterward, chromatographic separation was conducted under isocratic elution conditions using a Concise coregel 87H3 column (7.8 × 300 mm, 5 μm) (Concise Separations, CA, USA) and a mobile phase of 5 mM sulfuric acid, with the detection wavelength set at 210 nm. The other chromatographic conditions included a column oven temperature of 35 °C, a flow rate of 0.6 mL/min, and a run time of 60 minutes. Following chromatographic separation, individual SCFAs (acetic acid, propionic acid, and butyric acid) were identified and quantified based on the known standard retention times and peak areas (Sigma-Aldrich, MO, USA).

#### Pangenome analysis and strain specific gene extraction

After the *in vivo* experiment, the BPGA (Bacterial Pan Genome Analysis tool) v.1.3 computational pipeline was used to extract strain-specific genes for IDCC 2201 selected [19]. Seven annotated *S. thermophilus* genome sequences were used as input files. The unique genome database was constructed with a 50% identity cutoff using the USEARCH algorithm [20]. The strain-specific candidate gene for IDCC 2201 was confirmed to be specific for the strain analyzed through the basic local alignment search tool (BLAST). Genes with high target specificity were selected, and primers were designed using The PCR Primer Design Tool (Eurofins Genomics, Ebersberg, Germany) while considering amplicon size and GC content.

#### Strain-specific primer specificity and quantitative evaluation

Real-time PCR was performed using the CFX96 Deep Well Real-time System (Bio-Rad, CA, USA). The PCR mixture consisted of 20 ng of template DNA, 10 pmol/μL of primer pairs, 10 μL of 2×SYBR qPCR Mix (Toyobo, Osaka, Japan), and deionized distilled water to a total volume of 20 μL. Real-time PCR proceeded as follows: initial denaturation was initiated at 95 °C for 2 min, followed by forty cycles of 95 °C for 5 sec and 60 °C for 30 sec. A melting curve was obtained from 65°C to 95°C with holding and heating for 15 seconds at each step [21]. For the quantification of IDCC 2201 in fecal samples, a standard curve was constructed with the extracted IDCC 2201 genomic DNA. IDCC 2201 was cultured on MRS agar plates and counted after incubation at 37°C for 48 hours to confirm that the number of viable cells was 109 CFU/mL. Then, genomic DNA was extracted and diluted to 102 CFU/mL in deionized water using the serial decimal dilution method. Fecal samples were quantified by Log (CFU/g) based on the established standard curve.

#### Statistical analyses

All experimental data were presented as mean ± standard deviation. Differences between the control and experimental groups were analyzed by two-sample t-test or Kruskal–Wallis test to determine statistical significance. All statistical analyzes were performed in GraphPad Prism software (version 8.0). A between group p-value of 0.05 or less was considered to be representative of a statistically significant difference.

## Results & Discussion

### *In vitro* for screening probiotic candidate strains

#### Purine metabolism-related gene and analysis of hypoxanthine degrading

The *hpt* gene was identified in the *S. thermophilus* strain among the different types of strains according to the Ministry of Food and Drug Safety (MFDS, Cheongju, Republic of Korea). The *hpt* gene of *S. thermophilus* LMG 18311 was identified as UniProt ID: Q5M6K8, while the *hpt* gene of *S. thermophilus* CNRZ 1066 was identified as UniProt ID: Q5M216. The IDCC 2201 genome possessed the gene locus_tag=ESP48_05170 en-coding HGPRT. This protein is one of the essential enzymes in the purine salvage pathway that catalyzes the magnesium-dependent formation of IMP and GMP [22]. It is involved in the reduction of uric acid levels by recovering purines that are formed from the conversion of hypoxanthine and guanine to IMP and GMP [23,24]. This result suggests that IDCC 2201 can reduce uric acid levels through purine metabolism.

Considering the property described above, the ability of IDCC 2201 to degrade hypoxanthine was evaluated through HPLC. IDCC 2201 was shown to degrade 48.46% of 1.25 mM hypoxanthine for 1 hour. The development of strains that degrade purine compounds, such as hypoxanthine, has emerged as a potential treatment pathway for hyperuricemia [25]. It has been confirmed that there is a possibility of improving hyperuricemia by reducing uric acid through the inhibition of hypoxanthine.

#### Screening of LABs capable of degrading inosine

To select probiotics candidate strains, 15 candidate strains were evaluated for inosine degradation based on HPLC analysis. The degradation rate was calculated according to formula of degradation rate, and the results are shown in Fig 2. Among the 15 strains tested, the two strains with the highest inosine degradation rates were selected: 1.25 mM inosine was degraded to 84.65% inosine in *L. plantarum* CR1 and 99.98% inosine in *L. pentosus* GZ1. Inosine is an intermediate metabolite of purine metabolism that is related to the production of uric acid, which causes hyperuricemia [26]. The inosine degradation rate of strains selected in previous studies has been confirmed to range from 67.59% to 100% [16]. The inosine degrading abilities of the two strains selected in this study were 84.65% and 99.98%, which was confirmed to be similar to the values presented in a previous study. IDCC 2201, which has a gene related to purine metabolism and has been confirmed to have a hypoxanthine degrading ability, showed a degradation of 50.12% for inosine.

**Fig 2.**
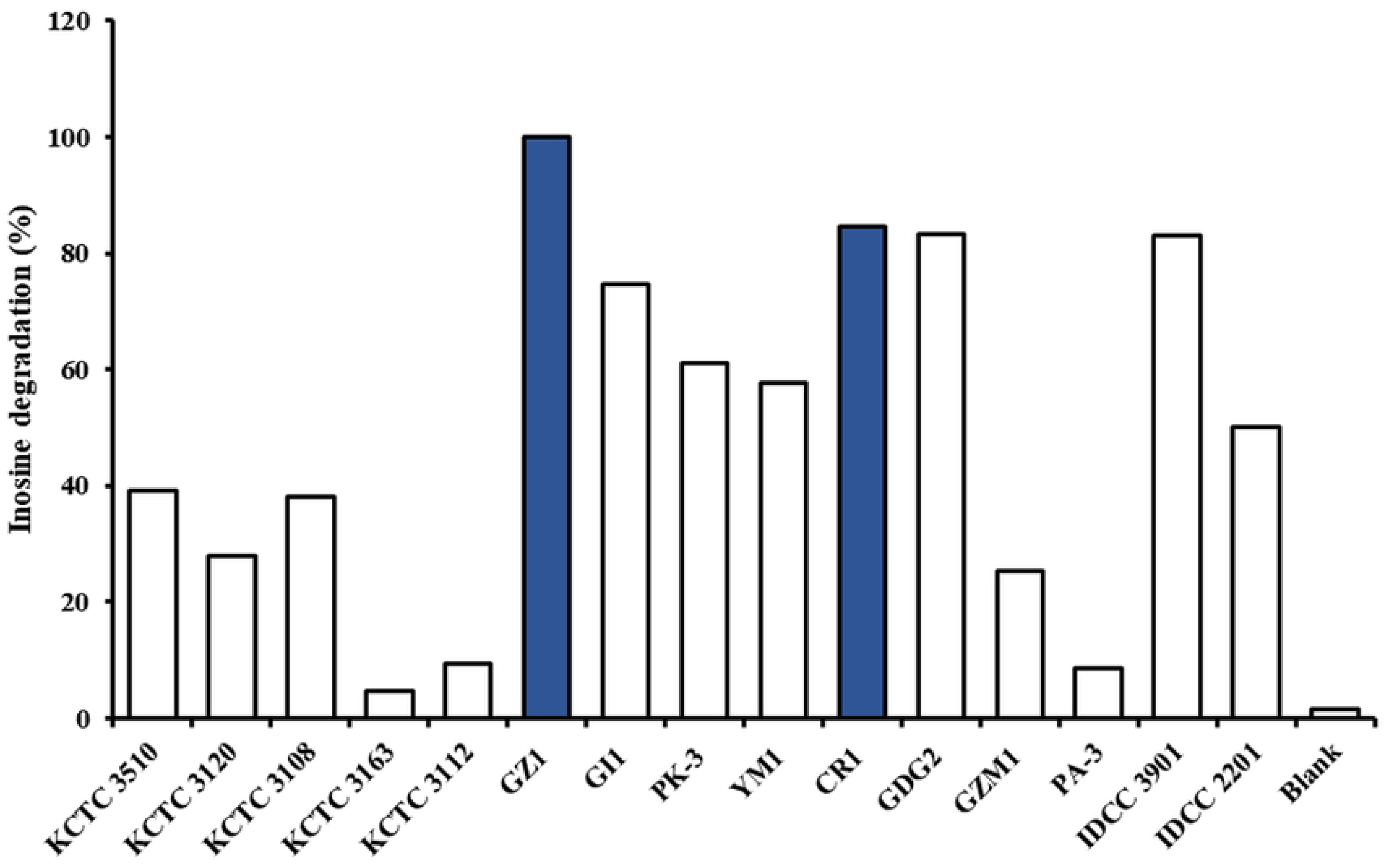
Inosine degradation analysis by HPLC.

By analyzing gene related to purine metabolism and confirming purine degradation, *L. plantarum* CR1, *L. pentosus* GZ1, and *S. thermophilus* IDCC 2201 strains were selected. Subsequent *in vivo* assays were performed using these three strains screened *in vitro*.

#### Efficacy evaluation of probiotics in rats with hyperuricemia

IDCC 2201, which was tested for hypoxanthine degradation, and CR1 and GZ1, which were tested for inosine degradation, all showed high degradation rates in the *in vitro* experiments. These three selected species were used in animal experiments that involved inducing hyperuricemia. SD rats with hyperuricemia induced by PO were used to study the effect of improving hyperuricemia [16]. PO, which in-creases blood uric acid level by inhibiting uricase, was used to examine the effect of candidate strains CR1, GZ1, and IDCC 2201 on blood uric acid level in hyperuricemia rats, and it was intraperitoneally administered for 7 days to induce hyperuricemia. LAB and allopurinol were orally administered 1 hour after PO administration for 7 days. During the experimental period, body weight increased in a statistically significant manner compared to G2 in all groups except for the G3 and G4 administration groups. The most important indicator of hyperuricemia is the uric acid content in the blood. As shown in Fig 3, the average level of uric acid in G2, which caused hyperuricemia, was 2.97 ± 1.01 mg/dL, which was statistically significantly increased by more than two times compared to the average level of 1.17 ± 0.15 mg/dL in G1. Mean-while, the uric acid levels were significantly decreased in the positive control groups, G8 and G9 (p < 0.05, p < 0.001). This indicates that the hyperuricemia model was successfully established.

**Fig 3.**
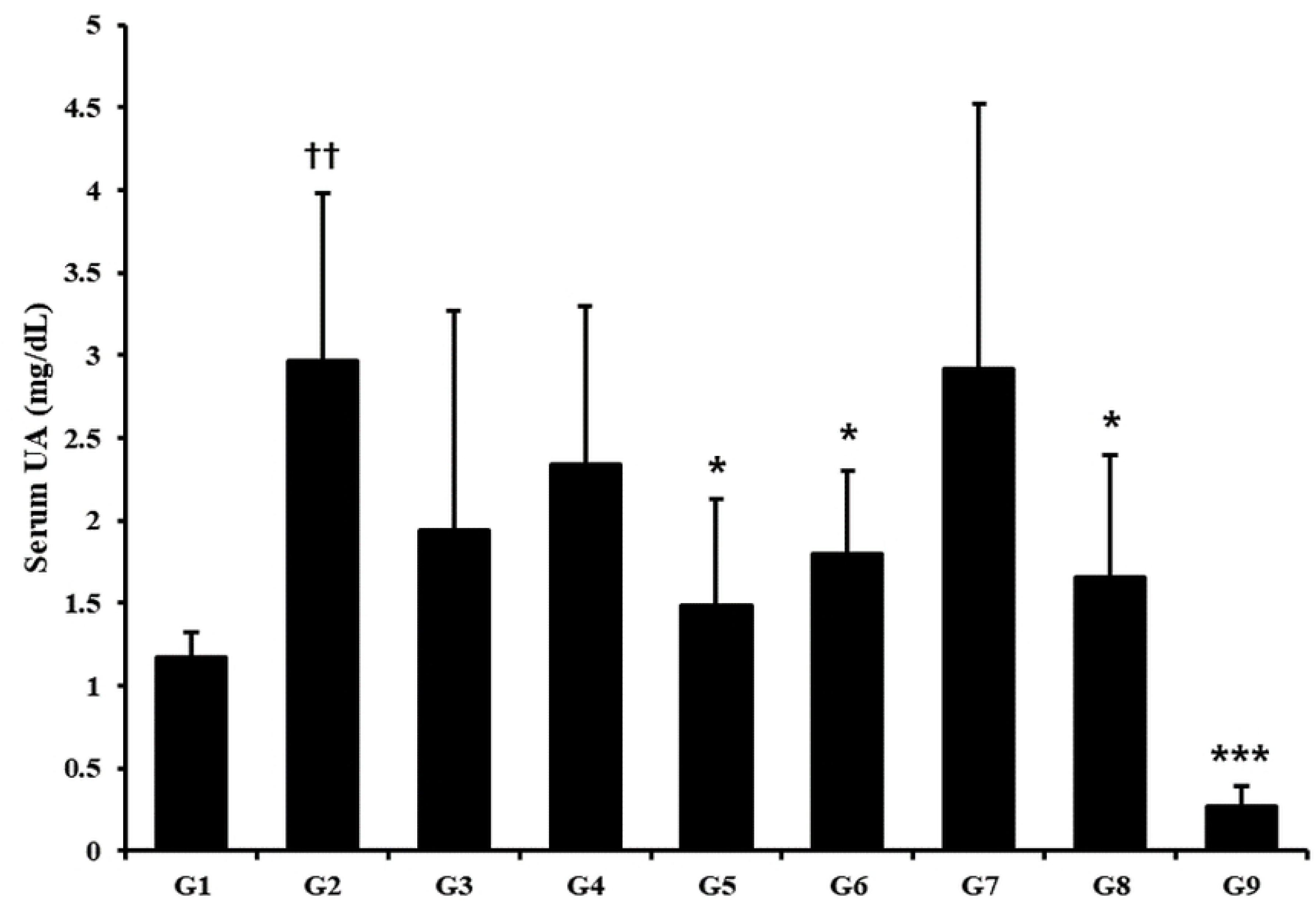
Effect of probiotics on the reduction of blood uric acid level in hyperuricemia rats. On the last day of administration of the test substance, blood was collected from the jugular vein of the rat 3 hours after administration of the test substance or the comparative substance, and the uric acid level was measured using a blood biochemical analyzer. G1: normal group; G2: hyperuricemia group; G3: CR1 (1×109 CFU/ day); G4: GZ1 (1×109 CFU/ day); G5: IDCC 2201 (1×109 CFU/ day); G6: CR1+IDCC 2201 (1×109 CFU/ day); G7: GZ1+IDCC 2201 (1×109 CFU/day); G8: PA-3 (1×109 CFU/ day); and G9: Allopurinol (50 mg/kg rat/day). Data are presented in the form of mean ± standard deviation values (N=6). Different symbols indicate significant differences according to statistical analysis through t-testing. †† p < 0.01 vs G1, * p < 0.05, *** p < 0.01 vs G2.

G5 and G6 significantly decreased uric acid levels by 1.48 ± 0.65 mg/dL and 1.8 ± 0.50 mg/dL, respectively (p < 0.05). Compared to the G2 group, the blood uric acid levels in the G5 and G6 administration groups were significantly reduced by 50.11% and 39.33%, respectively. These were effective values compared to those that were previously reported for *L. fermentum* NCUH003018 (30.77%) [16].

#### Effects of IDCC 2201 on the composition of gut microbiota in rats with hyperuricemia

Taxonomy analysis of the gut microbiota showed that PO-induced hyperuricemia treatment groups significantly altered the relative abundance of several bacterial phylum compared to G1. Compared to G1, hyperuricemia-induced G2 increased the relative abundance of Bacteroidetes, decreased the relative abundance of Firmicutes, and increased Actinobacteria. Treatment with IDCC 2201 increased the relative abundance of Firmicutes compared to G2, while treatment with allopurinol increased the relative abundance of Actinobacteria (Fig 4). As has previously been reported, IDCC 2201 reduced Bacteroidetes that were significantly increased by uric acid, and it increased Firmicutes that provide metabolic and immune properties to the host [27]. According to the analysis of the gut microbiome of gout patients, the ratio of Bacteroidetes to Firmicutes was higher in gout patients, and Bacteroidetes was also reported to be abundant in the gut microbiota of gout patients [28,29]. In a study analyzing the genes involved in uric acid degradation, it was reported that uric acid degradation was regulated by gut microbes, it was reported that Bacteroidetes possesses lipid A biosynthetic genes that contribute to inflammation, which may negatively affect endotoxin tolerance [29]. An imbalance in the ratio of Bacteroidetes/Firmicutes has also been observed in the gut microbiota of hyperuricemia mice [30]. It has been proven that achieving the control of the Bacteroidetes/Firmicutes ratio by these probiotics can improve hyperuricemia by controlling the intestinal microflora that has become unbalanced due to uric acid [31]. Actinobacteria was increased in G9 (allopurinol treated group), and a previous study reported that allopurinol treatment significantly affected the relative abundance of Actinobacteria [32]. The Shannon and Chao1 diversity indices were used to evaluate the alpha diversity of the gut microbiota. The results showed that allopurinol treatments significantly decreased both Shannon and Chao1 diversity indices compared to the IDCC 2201 and PO-induced hyperuricemia groups. PCA analysis based on unweighted UniFrac distance was conducted to evaluate the beta diversity of the gut microbiota. The results showed that IDCC 2201 induced a distinct clustering pattern compared to allopurinol treatment groups, thus indicating significant differences in gut microbiota composition. These findings suggest that IDCC 2201 treatment may have potential as a therapeutic strategy for hyperuricemia by modulating the gut microbiota. However, further studies are needed to elucidate the underlying mechanisms and confirm these results in human subjects.

**Fig 4.**
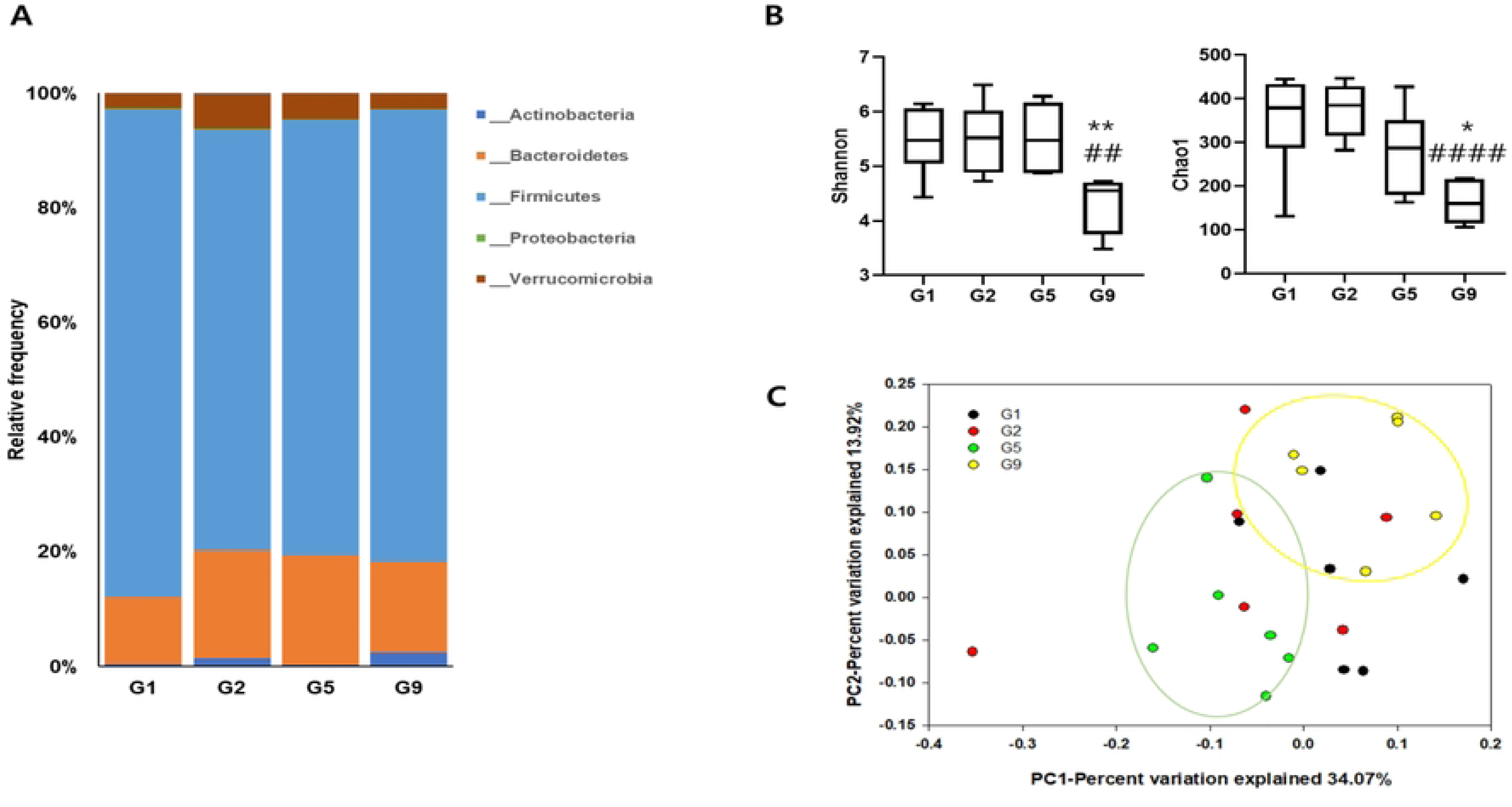
Effects of IDCC 2201 on gut microbiota in SD rats with PO-induced hyperuricemia. (A) Taxonomy analysis of community at phylum, (B) Shannon and Chao1 diversity, (C) PCA of unweighted UniFrac. G1: normal group; G2: hyperuricemia group; G5: IDCC 2201 (1×109 CFU/ day); and G9: Allopurinol (50 mg/kg rat/day). Data are expressed as mean ± standard deviation (n = 6). Compared with the G2 group, significance was determined as # p < 0.05, ## p < 0.01, ### p < 0.001, #### p < 0.0001, using the Kruskal–Wallis test. Compared with the G5 group, significance was determined as * p < 0.05, ** p < 0.01 using the Kruskal–Wallis test.

#### Uric acid reduction effect of IDCC 2201 by concentration in hyperuricemia rats

Based on the results derived from ‘Efficacy evaluation of probiotics in rats with hyperuricemia’, the reproducibility of uric acid reduction in the same animal model and dose dependency were evaluated for IDCC 2201 strain (Fig 5). There were no statistically significant differences between the weights of any test groups, and the average of G2 was 1.61 ± 0.42 mg/dL, which was a statistically significant increase compared to the average of 1.10 ± 0.20 mg/dL of G1 (p < 0.01). This indicates that an animal model has been successfully established. To confirm concentration dependence, IDCC 2201 was administered at concentrations of 1×109 CFU/day (IDCC 2201-SH, G3), 1×108 CFU/day (IDCC 2201-SM, G4), and 1×107 CFU/day (IDCC 2201-SL, G5), respectively. Group G3 was administered 1.18 ± 0.26 mg/dL, which was statistically significantly lower than that administered to G2 (p < 0.05). The G4 and G5 administration groups exhibited values of 1.45 ± 0.45 mg/dL and 1.38 ± 0.63 mg/dL, respectively, showing no significant difference from G2. This means that there is an effect of reducing uric acid in hyperuricemia when administered at a high concentration.

**Fig 5.**
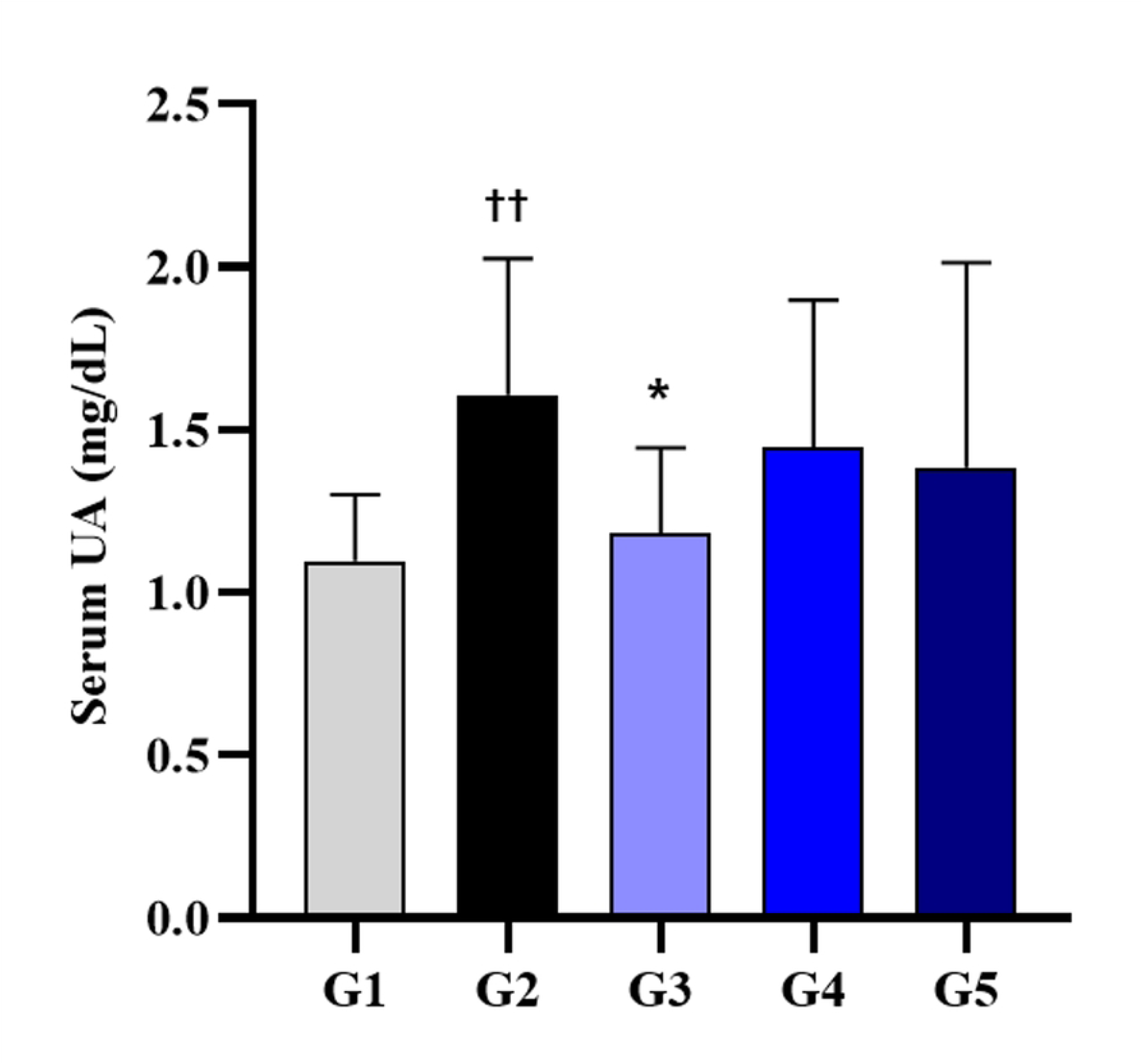
Effects of IDCC 2201 concentrations on the reduction of blood uric acid levels in hyperuricemia rats. On the 7th day of the test, 3 hours after administration of the test substance or the comparative substance, blood was collected from the animal’s jugular vein for the measurement of the uric acid level. G1: normal group; G2: hyperuricemia group; G3: IDCC 2201-SH (1×109 CFU/day); G4: IDCC 2201-SM (1×108 CFU/day); G5: IDCC 2201-SL (1×107 CFU/day). Data are presented in the form of mean ± standard deviation values (N=6). Different symbols indicate significant differences according to statistical analysis using t-testing. †† p < 0.01 vs G1, * p < 0.05 vs G2.

#### Analysis of the effect of IDCC 2201 on SCFAs in fecal samples

SCFAs were determined by HPLC in fecal samples of rats described in ‘Uric acid reduction effect of IDCC 2201 by concentration in hyperuricemia rats’. As shown in Fig 6, G2 showed significantly lower levels of both acetic acid (p < 0.01) and butyric acid (p < 0.001) compared to G1. On the other hand, G3 showed significantly higher levels of both acetic acid and butyric acid compared to G2 (p < 0.001, p < 0.001, respectively). In terms of the total content of SCFAs, there was a significant difference between G2 and G3 with p < 0.001.

**Fig 6.**
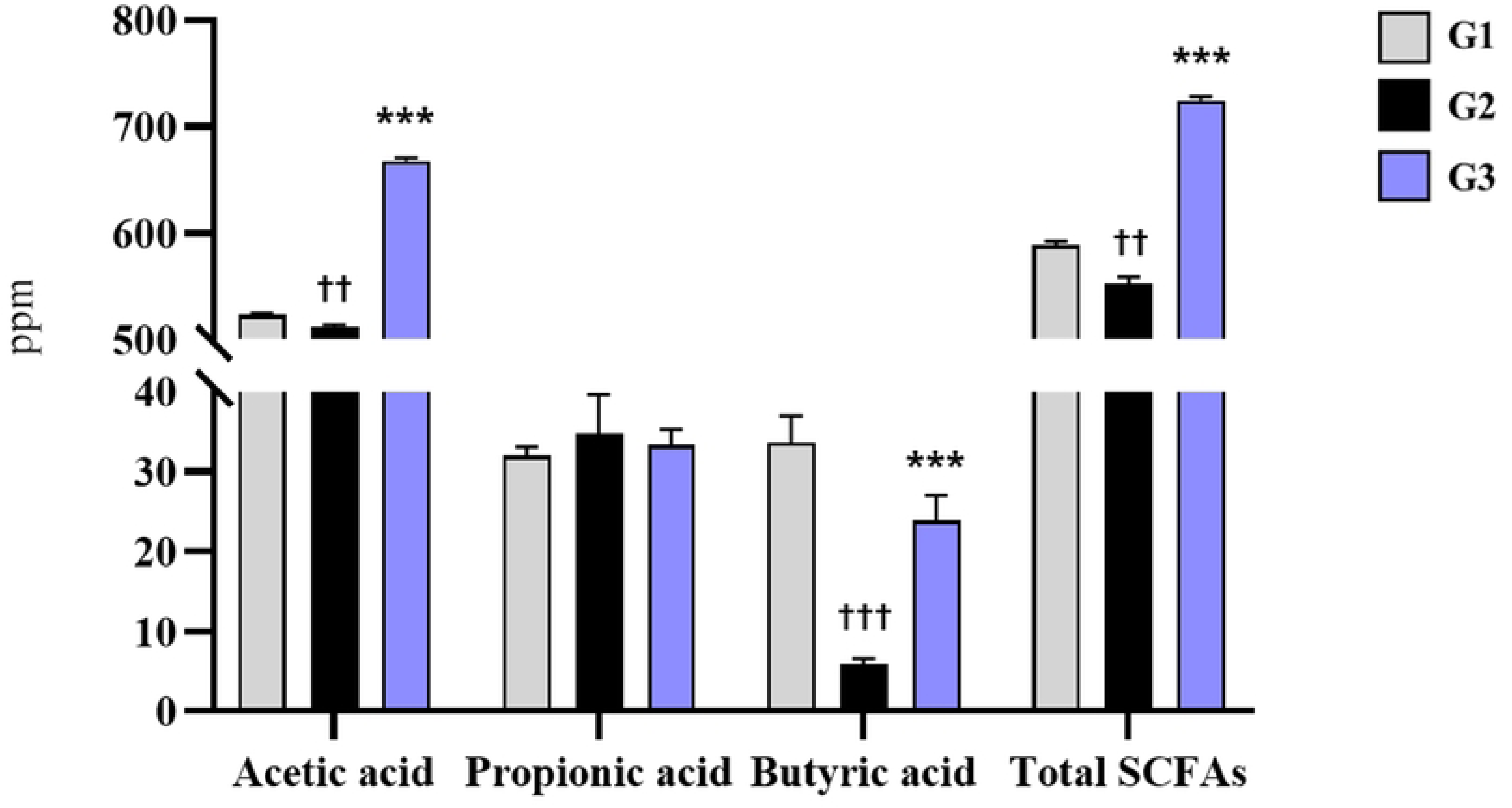
SCFAs content in rat feces. G1: normal group; G2: hyperuricemia group; G3: IDCC 2201-SH (1×109 CFU/day). The mean ± standard deviation values (N=6) are depicted by bars. Different symbols indicate significant differences according to statistical analysis using t-testing. †† p < 0.01, ††† p < 0.001 vs G1, *** p < 0.001 vs G2.

A previous study reported that hyperuricemia-induced rats have low intestinal bacterial abundance, thus resulting in low concentrations of SCFAs, and eventually leading to intestinal barrier dysfunction and increased intestinal permeability [33,34]. In a mouse model of hyperuricemia, SCFAs have been shown to be involved in uric acid metabolism as well as the intestinal barrier [35]. In particular, butyrate has been reported as having a beneficial effect on uric acid excretion by supplying ATP to the cells of the intestinal wall, thus improving uric acid metabolism and helping alleviate hyperuricemia [36,37]. Therefore, IDCC 2201 at a concentration of 1 × 109 CFU/day holds potential for helping reduce uric acid by showing an improvement effect on SCFAs that have been decreased due to hyperuricemia.

#### Analysis of IDCC 2201 using strain specific primer

With the 16S sequencing method, it is difficult to conduct quantitative evaluations at the bacterial species level [38]. To specifically detect the IDCC 2201 strain, real-time PCR analysis was performed using the flippase (Accession no. QAU28922.1) gene as a specific marker for IDCC 2201. IDCC 2201 in the fecal samples of G1, G2, and G3 was quantified with a standard curve (S1 Fig). The linearity range of the curve was 101–108 CFU/mL. The slope of the standard curve was −3.655 and the R2 value was 0.999, showing high efficiency [39]. It was identified at a significantly higher level in G3 compared to G1 and G2 (p < 0.001) (Fig 7). According to a previous study, after the administration of DM9218 (*Lactobacillus brevis*), which has the effect of improving hyperuricemia, *L. brevis* was detected in the feces of the DM9218-administered group, thus confirming a significant increase in the genus level [38]. In this study, IDCC 2201 was specifically detected at the strain level to verify whether IDCC 2201 delivered to the rat intestine was actually effective.

**Fig 7.**
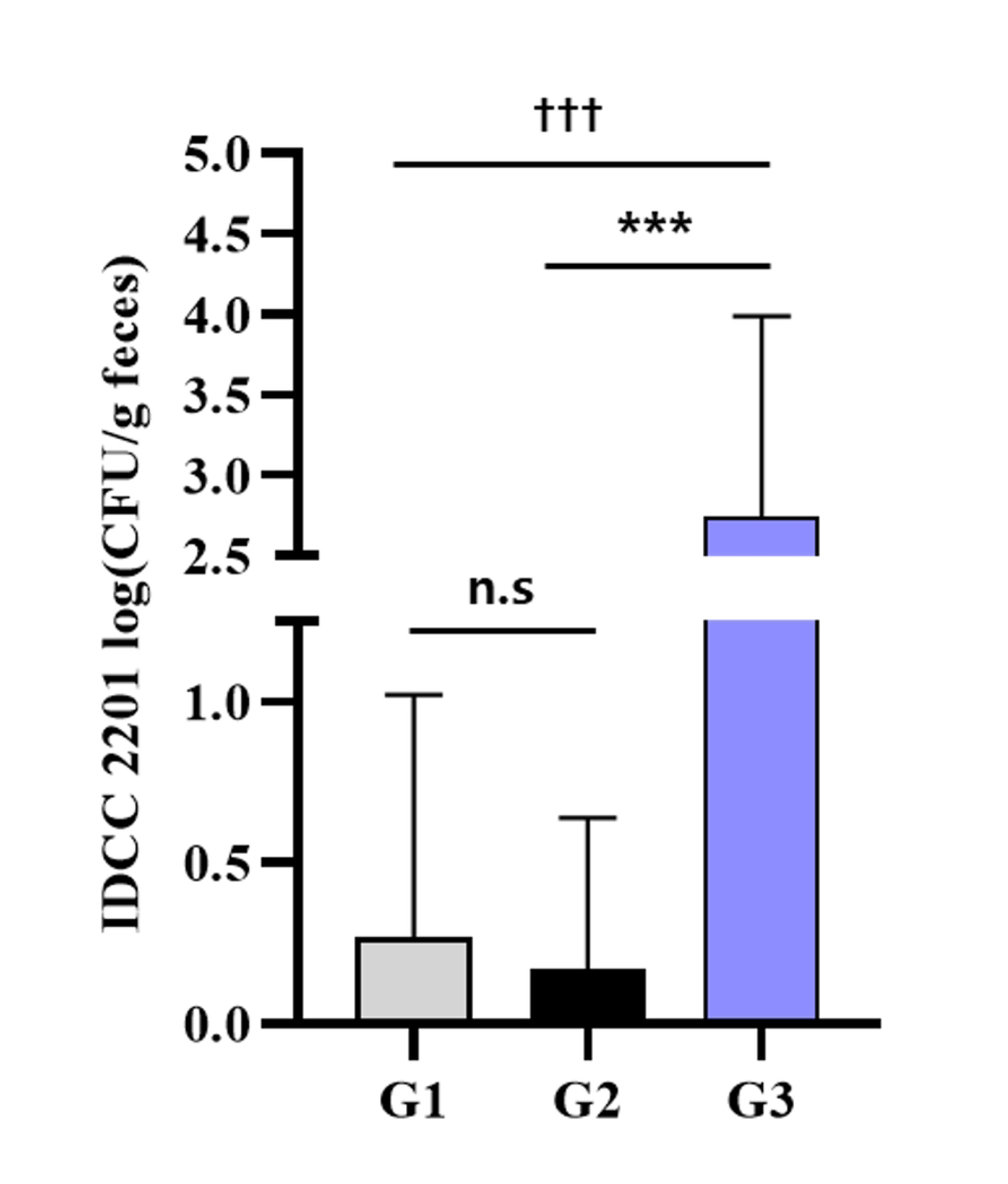
Quantification of IDCC 2201 in fecal samples of rats with hyperuricemia administered with IDCC 2201, as analyzed by real-time PCR. G1: normal group; G2: hyperuricemia group; G3: IDCC 2201-SH (1×109 CFU/day). Data are presented in the form of mean ± standard deviation values (N=6). Different symbols indicate significant differences according to statistical analysis using t-testing. n.s: no significant difference between groups. ††† < 0.001 vs G1, *** p < 0.001 vs G2.

### Conclusion

This study evaluated the possession of gene related to purine metabolism and the ability to degrade purines, and a probiotics strain that was able to both reduce blood uric acid levels and improve the intestinal environment was identified through animal experiments involving the induction of hyperuricemia. The selected IDCC 2201 possesses an enzyme-related gene that decomposes hypoxanthine, and it was found to exhibit inosine and hypoxanthine degrading ability *in vitro*. Moreover, the serum uric acid concentration of hyperuricemia rats was improved, and reproducibility was confirmed at a concentration of 1 × 109 CFU/day. Further, IDCC 2201 lowered the ratio of Bacteroidetes/Firmicutes, increased biodiversity, and significantly increased SCFAs in the intestine. This was demonstrated to be ameliorated by an increase in IDCC 2201 in fecal samples, thus showing the uric acid reducing and intestinal health functions of IDCC 2201. Given the critical role of gut microbiome composition in human health and disease through interactions between gut microbiota and human health, probiotics could be a promising strategy when used as living therapeutics to prevent or treat disease [40]. There is a need for metabolic studies to elucidate the mechanism by which IDCC 2201 prevents hyperuricemia, and additional verification through clinical trials is needed to continue developing it as a practical treatment.

## Supporting information

**S1 Fig. Real-time PCR standard curve for IDCC 2201.**

